# Regional and tissue-specific metabolic differences in human neural retina and RPE/choroid

**DOI:** 10.64898/2026.08.04.742646

**Authors:** Ting Zhang, Yinxiao Xiang, Mark C. Gillies, Ling Zhu, Jianhai Du

## Abstract

It is clear that the human retina and its underlying retinal pigment epithelium and choroid (RPE/choroid) form an interdependent metabolic ecosystem, but how metabolism differs between the cone-rich macula and rod-rich periphery remains unclear. Using targeted metabolomics, we quantified 133 metabolites in paired macular and peripheral neural retina and RPE/choroid explants from human donor eyes following short-term culture to restore metabolic activity. Distinct metabolic differences were identified between retinal regions and between tissues. Compared with the peripheral retina, the macula showed metabolic features consistent with greater glycolytic activity, increased NADH availability and higher levels of the neurotransmitter-associated metabolites N-acetyl-aspartate (NAA) and N-acetyl-aspartyl-glutamate (NAAG), consistent with increased energetic and neuronal activity. Compared with peripheral RPE/choroid, the macular RPE/choroid had higher levels of the flavin cofactor FAD together with NAD-related metabolites, including NAD, NADP and NAAD. Comparisons between the neural retina and RPE/choroid further showed that the neural retina was primarily associated with energy production and neurotransmission, whereas the RPE/choroid was associated with cofactor metabolism, nucleotide salvage and lipid metabolism. These findings are consistent with metabolic coupling between the neural retina and RPE/choroid. The macula has metabolic features consistent with high energetic demand, providing a potential metabolic basis for its selective vulnerability in macular disease.

## Introduction

The human retina is one of the most energy-demanding tissues in the body (1). Its high metabolic demand is supported by close coordination between the neural retina, retinal pigment epithelium (RPE) and choroid, which form an interdependent metabolic ecosystem. The RPE facilitates nutrient delivery, metabolic waste removal, light absorption and intermediate recycling, while the choroid provides the blood supply essential for oxygen and nutrient transport (2, 3). Disruption of this metabolic cooperation is associated with major retinal diseases including age-related macular degeneration (AMD) and diabetic retinopathy, in which mitochondrial dysfunction, altered lipid metabolism and reduced choroidal perfusion have been implicated in disease pathogenesis (4–9).

The RPE and neural retina are metabolically coupled to maintain retinal homeostasis and function. We have previously reported that the neural retina requires substantial uptake of glutamate, aspartate and other amino acids to support carbon and nitrogen metabolism (10). The RPE preferentially uses proline and converts it into mitochondrial intermediates and amino acids including citrate, glutamate and aspartate, which are exported to the neural retina to support retinal metabolism (11). The neural retina produces energy by converting glucose to lactate, which the RPE uses as a direct energy source, helping preserve glucose for the neural retina (12, 13). By phagocytosing shed photoreceptor outer segments, the RPE recycles lipids that can be reused to support photoreceptor renewal and neural retinal homeostasis (2).

The macula, responsible for central vision, is structurally and functionally distinct from the peripheral retina. It has a higher density and diversity of retinal cells, particularly cone photoreceptors and ganglion cells, whereas the peripheral retina is enriched in rod photoreceptors (14, 15). The RPE also differs between the macula and the periphery. In the macula, RPE cells are smaller, more densely packed, more uniform and more pigmented while the peripheral RPE cells are larger and less pigmented (16, 17). Such structural and functional features may be associated with regional metabolic specialisation. We have observed regional differences in gene expression, as well as in metabolite uptake from and release into the culture medium, in both neural retinal and RPE/choroid explants from macula and periphery (10). However, understanding metabolic interactions between human retinal and RPE/choroid tissues under physiological conditions remains a major challenge because postmortem tissues are affected by ischemic changes and cellular deterioration.

To better capture metabolically active tissue states, we cultured paired human macular and peripheral explants, including neural retina and RPE/choroid, in nutrient-rich medium for four hours, allowing partial restoration of metabolic activity. We then quantified tissue metabolites using targeted liquid chromatography-mass spectrometry (LC-MS). This approach allowed us to compare macular and peripheral regions, as well as neural retina and RPE/choroid within each region and characterise regional and tissue-specific metabolic differences in the human retina and RPE/choroid.

## Results

### Metabolic differences between human macula and peripheral neural retina

Paired macular and peripheral retinal explants were cultured and collected for targeted metabolomic analysis (**Fig. 1A**). Partial least squares-discriminant analysis (PLS-DA) of all detected metabolites showed moderate but distinct separation between macular and peripheral neural retina samples (**Fig. 1B**). The first two PLS-DA components showed separation between the groups, with component 1 and component 2 accounting for 34.6% and 41.1%, respectively. Volcano plot analysis identified 43 significantly different metabolites, of which 9 were higher in the macula and 34 were higher in the peripheral retina (**Fig. 1C**).

**Figure 1.**
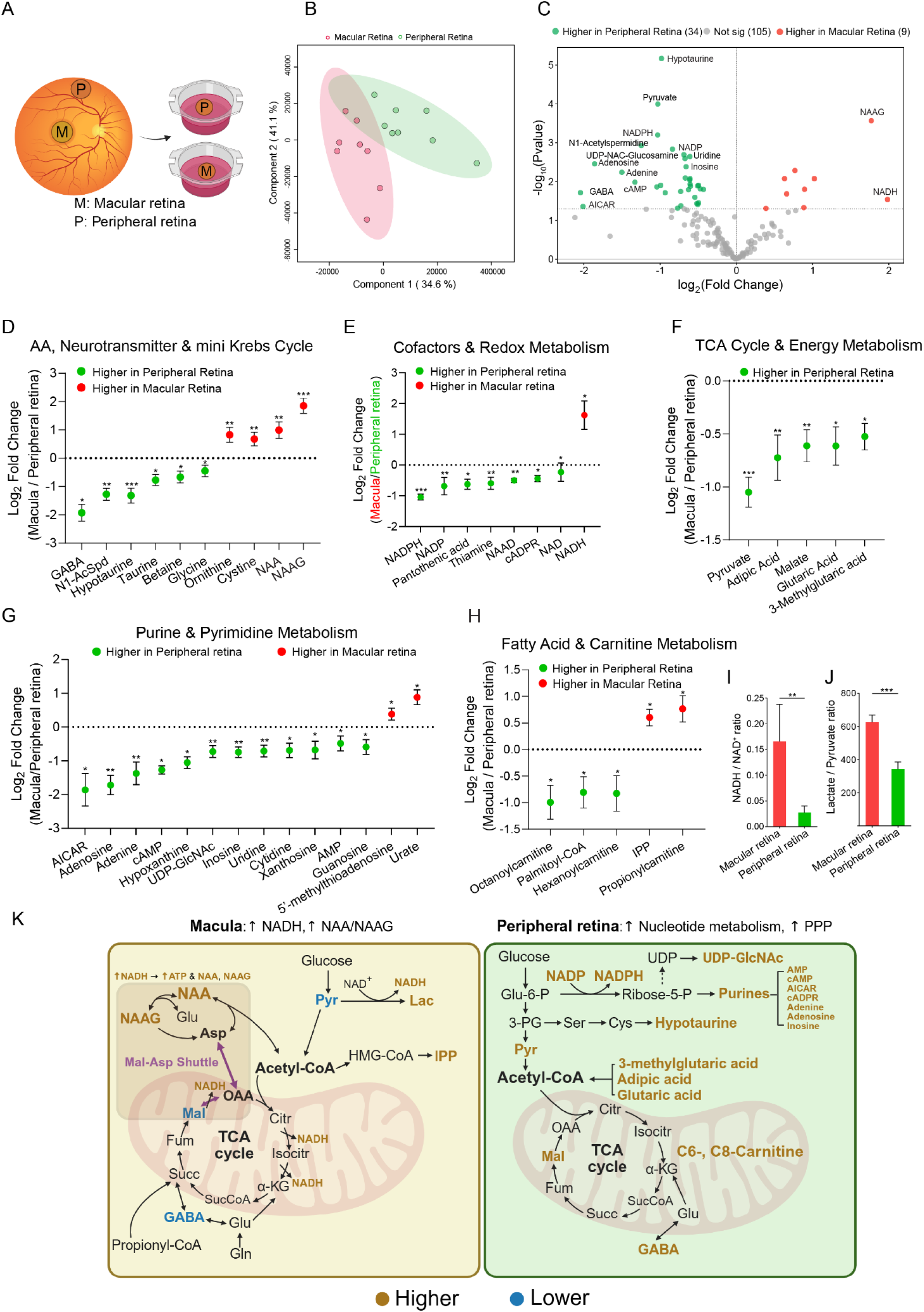
Metabolic differences between the macula and peripheral neural retina. **A**. Schematic of human neural macula (M) and peripheral (P) retina cultured on transwell inserts for four hours. **B**. PLS-DA showing distinct clusters of metabolite profiles between the macula (red) and peripheral retina (green). **C**. Volcano plot showing significantly different metabolites between the macula and peripheral retina. Red dots indicate higher levels in the macula and green dots indicate higher levels in the periphery. **D–H**. Log₂ fold-change plots (macula relative to periphery) showing differential metabolite levels across: **D**. amino acids, neurotransmitters and mini-Krebs cycle intermediates; **E**. cofactors and redox metabolism; **F.** TCA cycle and energy metabolism; **G**. purine and pyrimidine metabolism; **H**. fatty acid and carnitine metabolism. **I**. NADH/NAD⁺ ratio in macula versus periphery. **J**. Lactate/pyruvate ratio in macula versus periphery. **K**. Schematic summary of regional metabolic differences. “Higher” and “Lower” indicate metabolites with higher or lower abundance, respectively, in the macular or peripheral neural retina relative to the other region. Error bars represent standard error across biological replicates. n=8 donors. *p < 0.05, **p < 0.01 and ***p < 0.001.

Neurotransmitter-related metabolites were significantly higher in the macula, particularly N-acetyl-aspartate (NAA) and N-acetyl-aspartyl-glutamate (NAAG) (**Fig. 1D**). The macula was also enriched in antioxidant-related metabolites (urate, cystine) and lipid-related intermediates (isopentenyl pyrophosphate [IPP], propionylcarnitine) (**Fig. 1D, G-H**). The macula had higher NADH/NAD⁺ and lactate/pyruvate ratios, consistent with increased glycolytic and NADH-linked energy metabolism (**Fig. 1I-J**).

Compared with the neural macula, the peripheral retina had higher levels of metabolites in purine metabolism (AMP, adenosine, inosine, hypoxanthine) and the pentose phosphate pathway (NADPH, NADP⁺, thiamine) (**Fig. 1E, G**). Pantothenic acid, a cofactor related to coenzyme A synthesis, was also higher in the peripheral retina (**Fig. 1E**). In addition, the peripheral retina had higher levels of taurine, betaine and γ-aminobutyric acid (GABA), which are involved in osmolyte and neurotransmitter metabolism, together with fatty acid β-oxidation-related metabolites, including hexanoylcarnitine (C6-carnitine) and octanoylcarnitine (C8-carnitine) (**Fig. 1D, H)**.

These findings highlight distinct regional metabolic profiles between the macular and peripheral neural retina. The neural macula showed higher glycolysis and NADH-related features, whereas the peripheral retina had higher levels of metabolites associated with purine metabolism, the pentose phosphate pathway and medium-chain acylcarnitines. These regional metabolic differences are summarised schematically in **Fig. 1K**.

### Metabolic differences between the macular and peripheral RPE/choroid

Paired macular RPE/choroid and peripheral RPE/choroid explants were cultured and collected for targeted metabolomic analysis (**Fig. 2A**). PLS-DA of all detected metabolites found a moderate separation between the macular and peripheral RPE/choroid (**Fig. 2B**). The first two components accounted for most of the variation in the data, with component 1 explaining 29.6% and component 2 explaining 49.5% of the total variance. Volcano plot analysis identified 12 significantly different metabolites, with 7 higher in the macular RPE/choroid and 5 higher in the peripheral RPE/choroid (**Fig. 2C**).

**Figure 2.**
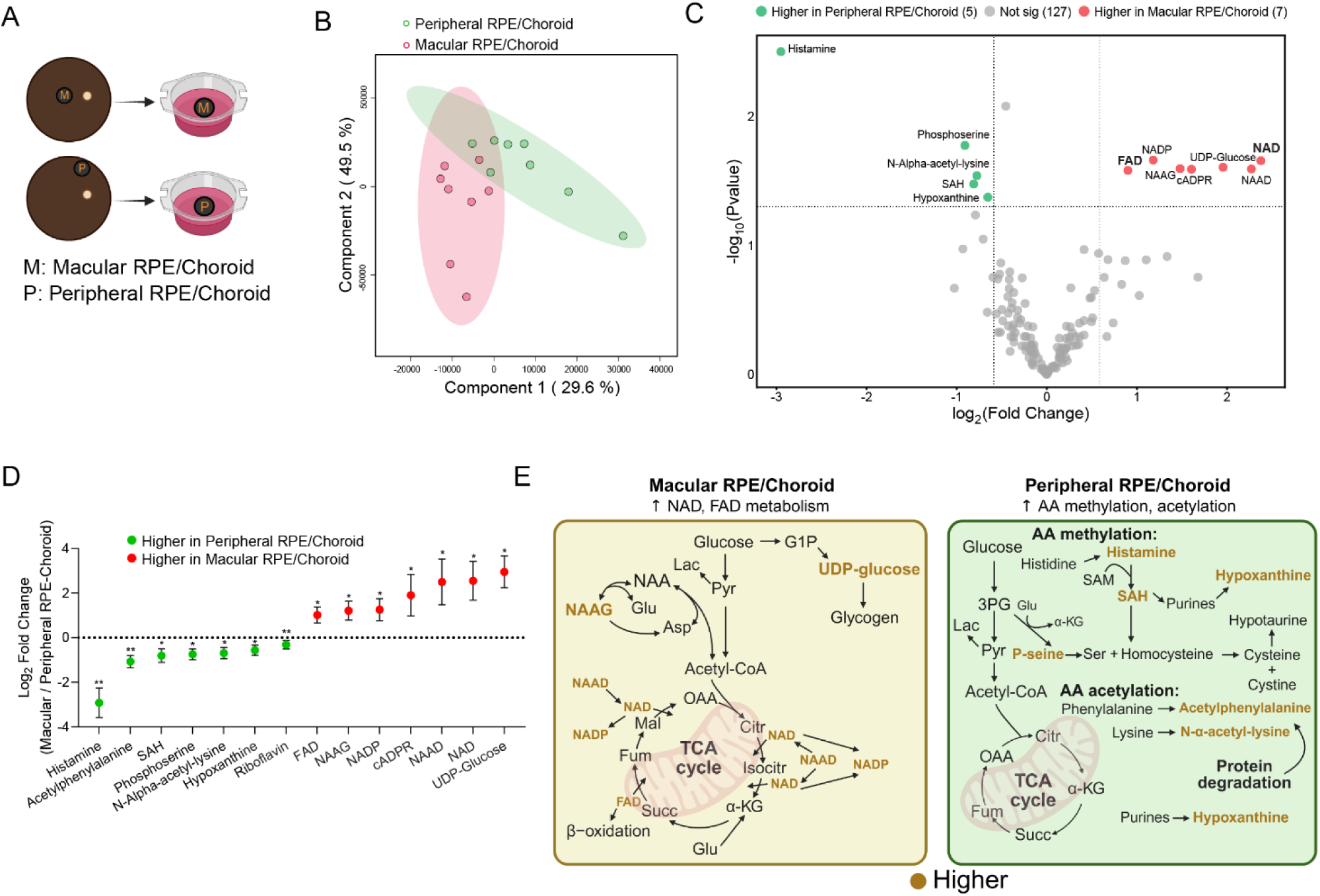
Metabolic differences between the macular and peripheral RPE/choroid. **A**. Schematic of human macular and peripheral RPE/choroid cultured on transwell inserts for four hours. **B**. PLS-DA showing separation of the macular (red) and peripheral RPE/choroid (green) based on metabolite profiles. **C**. Volcano plot showing metabolites with significantly higher levels in the macular RPE/choroid (red) or peripheral RPE/choroid (green). Key metabolites are annotated. **D**. Log₂ fold changes of metabolites in the macular RPE/choroid relative to the peripheral RPE/choroid. **E**. Schematic summary of regional metabolic differences. Brown indicates metabolites present at higher levels in the macular or peripheral RPE/choroid relative to the other region.

Log₂ fold-change analysis showed that the macular RPE/choroid had higher levels of the flavin cofactor FAD and NAD-related metabolites, including NAD, NADP, NAAD and cADPR, as well as the neurotransmitter-associated metabolite NAAG and the nucleotide sugar UDP-glucose, than peripheral RPE/choroid explants (**Fig. 2D**). Peripheral RPE/choroid had higher levels of amino acid-related metabolites, including acetylphenylalanine, N-alpha-acetyl-lysine, phosphoserine and histamine, together with the methylation-related metabolite S-adenosylhomocysteine (SAH) and the purine salvage intermediate hypoxanthine (**Fig. 2D**).

These findings again indicate regional metabolic differences between the macular and peripheral RPE/choroid, with more NAD-related cofactors in the macula and more amino acid modification, methylation-related and purine salvage metabolites in the periphery. A schematic summary of these metabolic differences is shown in **Fig. 2E**, with metabolites higher in each region highlighted.

### Distinct metabolic profiles in human neural macula and macular RPE/choroid

Paired neural macula and macular RPE/choroid explants were cultured and collected for targeted metabolomic analysis (**Fig. 3A**). PLS-DA of all detected metabolites showed clear separation between neural macula and macular RPE/choroid samples (**Fig. 3B)**, indicating distinct metabolic profiles between the two tissues. The first two components accounted for 28.2% and 4.9% of the variation, respectively. The volcano plot identified 30 metabolites that were significantly higher in the neural macula and 27 metabolites that were significantly higher in the macular RPE/choroid (**Fig. 3C**).

**Figure 3.**
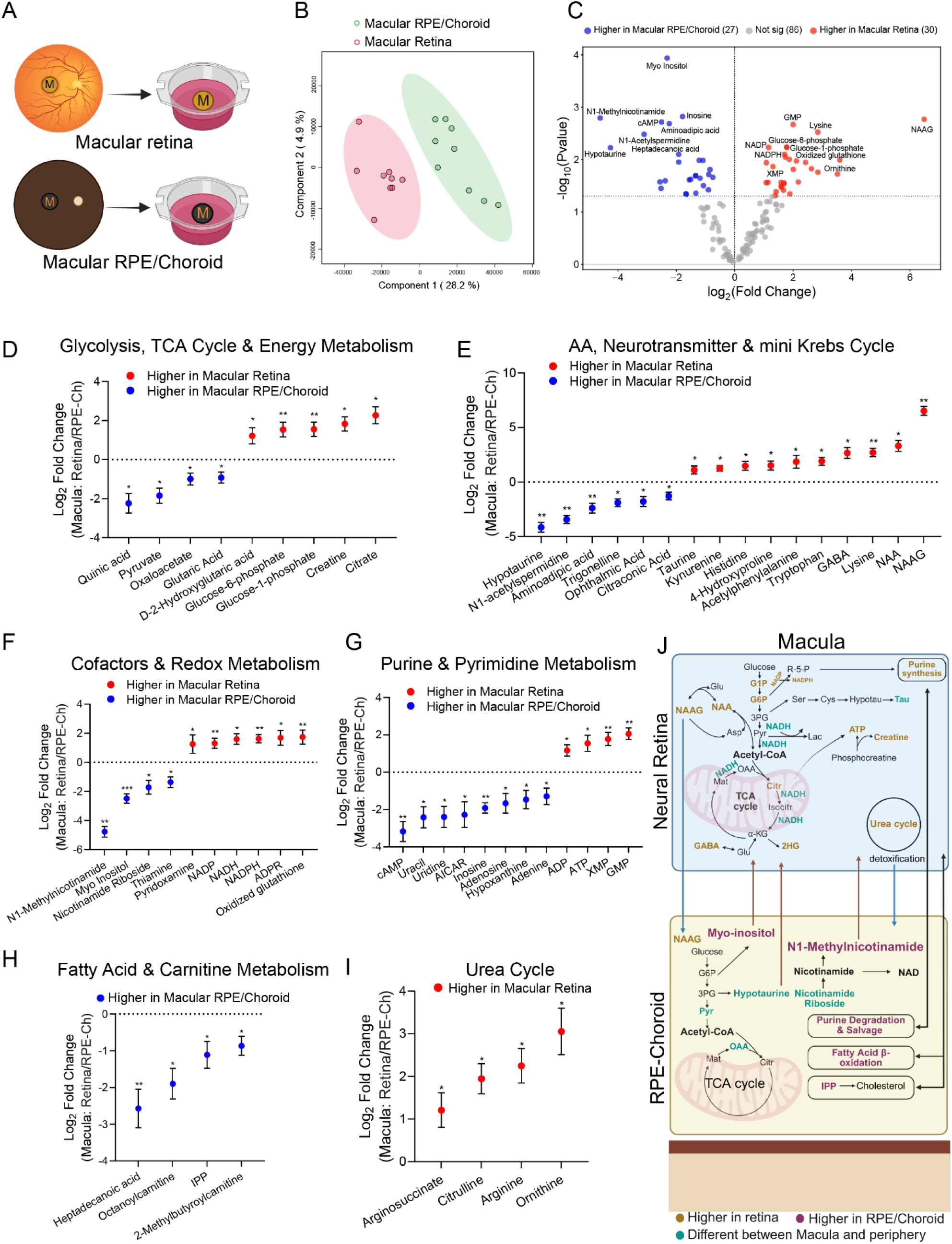
Comparative metabolic profiling of neural macula and macular RPE/choroid. **A**. Schematic of human neural macula and RPE/choroid cultured on transwell inserts for four hours. **B**. PLS-DA showing separation of the neural macula (red) and macular RPE/choroid (green) based on metabolite profiles. **C**. Volcano plot showing metabolites with significantly higher levels in the neural macula (red) or the macular RPE/choroid (blue). Key metabolites are annotated. **D–I**. Log₂ fold changes of metabolites (macular retina relative to macular RPE/choroid) grouped by pathway: glycolysis/TCA cycle & energy metabolism (**D**), amino acid, neurotransmitter & mini-Krebs cycle metabolites (**E**), cofactors & redox metabolism (**F**), purine & pyrimidine metabolism (**G**), fatty acid & carnitine metabolism (**H**) and urea cycle (**I**). **J**. Schematic summary of metabolic pathways in the human macula. Brown indicates metabolites with higher abundance in the neural macula; purple indicates metabolites with higher abundance in the macular RPE/choroid; green highlights metabolites that differed between the macula and periphery within the same tissue.

Compared with the macular RPE/choroid, the neural macula had higher levels of glycolytic and TCA cycle metabolites, including glucose-6-phosphate, glucose-1-phosphate and citrate, together with the energy-related metabolite creatine (**Fig. 3D**). Amino acids and amino acid derivatives, including histidine, lysine and tryptophan, were also higher, together with neurotransmitter-related metabolites such as GABA, taurine, kynurenine, NAA and NAAG (**Fig. 3E**). Metabolites involved in redox metabolism, including pyridoxamine, NADP, NADPH and oxidised glutathione, were also higher in the neural macula (**Fig. 3F**). Purine nucleotides, including ATP, ADP, GMP and XMP, were enriched in the neural macula (**Fig. 3G**). In addition, urea cycle intermediates, including citrulline, arginine, ornithine and argininosuccinate, were more abundant in the neural macula (**Fig. 3I**).

The macular RPE/choroid had higher levels of organic acids and related metabolites, including glutaric acid, oxaloacetate and quinic acid, than the neural macula (**Fig. 3D**). Amino acid-related metabolites, including aminoadipic acid, hypotaurine, ophthalmic acid and N1-acetylspermidine, were also higher (**Fig. 3E**). Vitamin B-derived and NAD-related metabolites, including thiamine, nicotinamide riboside and N1-methylnicotinamide, were enriched in the macular RPE/choroid, with N1-methylnicotinamide approximately 16-fold higher (**Fig. 3F**). Nucleosides and related metabolites, including inosine, adenosine, uridine and cAMP, were higher in the macular RPE/choroid (**Fig. 3G)**. Lipid-related metabolites, including acylcarnitines and IPP, were also higher in the macular RPE/choroid (**Fig. 3H**).

The schematic summary highlights the main metabolic differences between the neural macula and macular RPE/choroid, with retina-enriched metabolites shown in brown and RPE/choroid-enriched metabolites shown in purple (**Fig. 3J**).

### Distinct metabolic profiles in human peripheral neural retina and peripheral RPE/choroid

Paired peripheral neural retina and peripheral RPE/choroid explants were cultured and collected for targeted metabolomic analysis (**Fig. 4A**). PLS-DA of all detected metabolites had a clear separation between peripheral neural retina and peripheral RPE/choroid samples, indicating distinct metabolic profiles between the two tissues (**Fig. 4B**). Component 1 accounted for 35.8% of the variation, while component 2 accounted for 30.3%. The volcano plot had 32 metabolites with significantly higher levels in the peripheral retina and 15 metabolites with significantly higher levels in the peripheral RPE/choroid (**Fig. 4C**).

**Figure 4.**
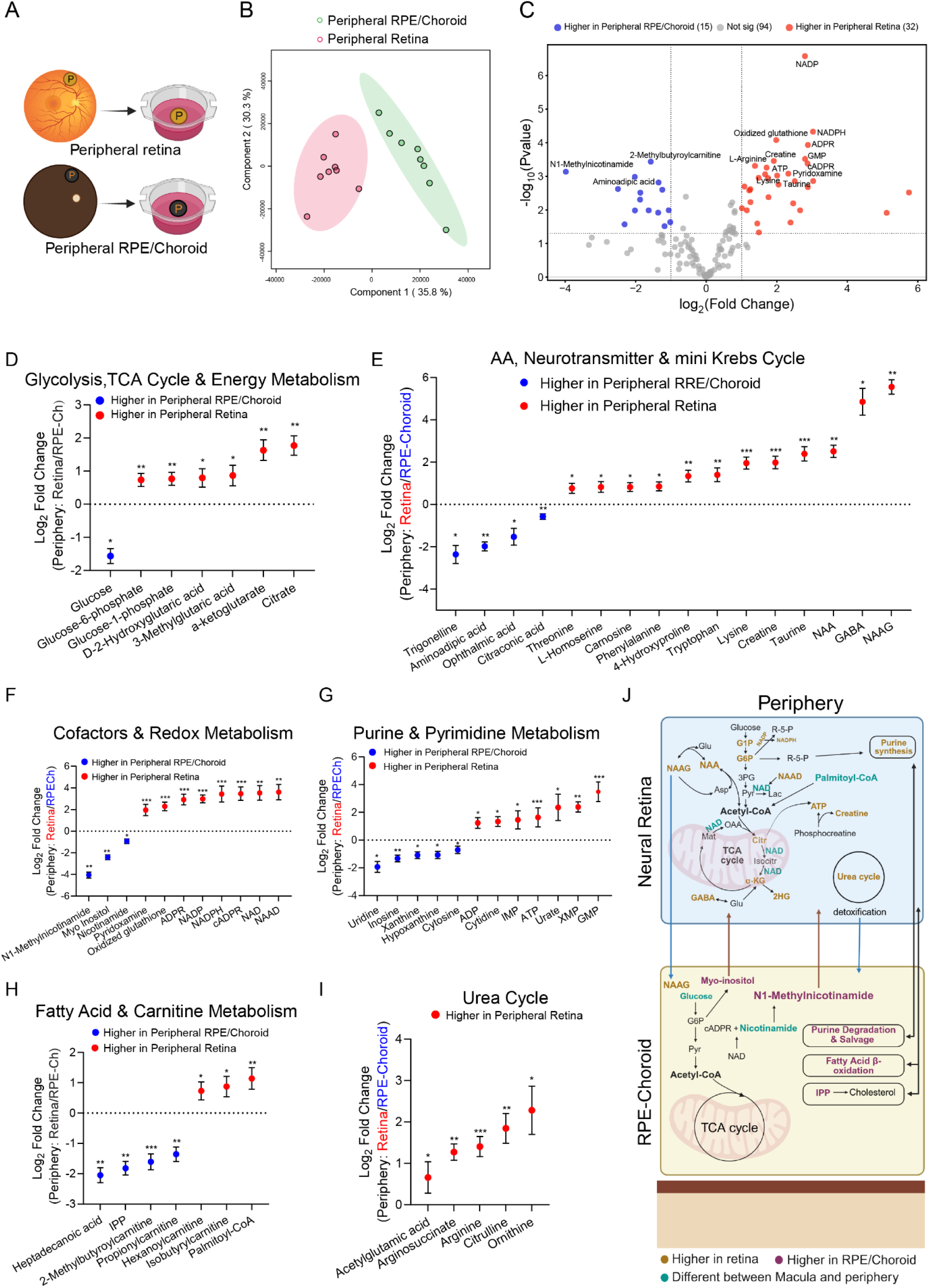
Comparative metabolic profiling of peripheral retina and peripheral RPE/choroid. **A**. Schematic of human peripheral retina and RPE/choroid cultured on transwell inserts for four hours. **B**. PLS-DA showing separation of the peripheral retina (red) and peripheral RPE/choroid (green) based on metabolite profiles. **C**. Volcano plot showing metabolites significantly higher in the peripheral retina (red) or in the peripheral RPE/choroid (blue). **D**–**I**. Log₂ fold changes of metabolites (peripheral retina relative to peripheral RPE/choroid) grouped by pathway: glycolysis/TCA cycle & energy metabolism (**D**), amino acid, neurotransmitter & mini-Krebs cycle metabolites (**E**), cofactors & redox metabolism (**F**), purine & pyrimidine metabolism (**G**), fatty acid & carnitine metabolism (**H**) and urea cycle (**I**). **J.** Schematic summary of metabolic pathways in the human peripheral retina and RPE/choroid. Brown indicates metabolites with higher abundance in the peripheral neural retina; purple indicates metabolites with higher abundance in the peripheral RPE/choroid; green highlights metabolites that differed between the macula and periphery within the same tissue.

Compared with the peripheral RPE/choroid, the peripheral retina had higher levels of glycolytic and TCA cycle metabolites, including glucose-6-phosphate, citrate and α-ketoglutarate (**Fig. 4D**). Amino acids and neurotransmitter-related metabolites, including tryptophan, taurine, GABA, NAA and NAAG, were also enriched in the peripheral retina (**Fig. 4E**). Metabolites involved in redox metabolism, including NADP, NADPH and oxidised glutathione, were higher (**Fig. 4F**). Purine and pyrimidine metabolites, including ATP, GMP and XMP, were also higher in the peripheral retina (**Fig. 4G**). In addition, urea cycle intermediates, including arginine, citrulline and ornithine, together with lipid-related metabolites such as palmitoyl-CoA, were more abundant in the peripheral retina (**Fig. 4H–I**).

The peripheral RPE/choroid had higher levels of amino acid-related metabolites, including aminoadipic acid and ophthalmic acid (**Fig. 4E**). Metabolites related to NAD metabolism, including nicotinamide and N1-methylnicotinamide, were enriched in the peripheral RPE/choroid (**Fig. 4F**). Purine and pyrimidine metabolites, including inosine and hypoxanthine, were also higher (**Fig. 4G**). In addition, lipid-related metabolites, including IPP and acylcarnitines, were enriched in the peripheral RPE/choroid (**Fig. 4H**).

The schematic summary highlights the main metabolic differences between the peripheral neural retina and peripheral RPE/choroid, with retina-enriched metabolites shown in brown and RPE/choroid-enriched metabolites shown in purple (**Fig. 4J**).

### Distinct metabolic patterns across retinal regions and tissues

Heatmap analysis of averaged metabolite levels was used to visualise overall metabolic trends across the macular and peripheral neural retina and RPE/choroid, showing broad tissue-specific and regional metabolic patterns (**Fig. 5**). Overall, the neural retina had a general enrichment of amino acids and amino acid derivatives, particularly in the neural macula (**Fig. 5A**). Energy-related metabolites, including glycolytic intermediates and creatine also tended to be higher in the neural retina. Several downstream TCA cycle intermediates had relatively higher levels in the RPE/choroid (**Fig. 5B**). Lipid- and fatty acid-related metabolites had distinct tissue-dependent patterns. Some acylcarnitines and fatty acid metabolism-associated metabolites were relatively enriched in the RPE/choroid, whereas palmitoyl-CoA and some carnitine-related metabolites were higher in the neural retina (**Fig. 5C**). Nucleotide and nucleoside metabolites also had clear tissue-specific distributions. ATP, ADP, GMP and XMP tended to be higher in the neural retina, whereas inosine, adenosine, uridine and related metabolites were more abundant in the RPE/choroid (**Fig. 5D**). Vitamin, cofactor and redox-related metabolites had distinct patterns between tissues.

**Figure 5.**
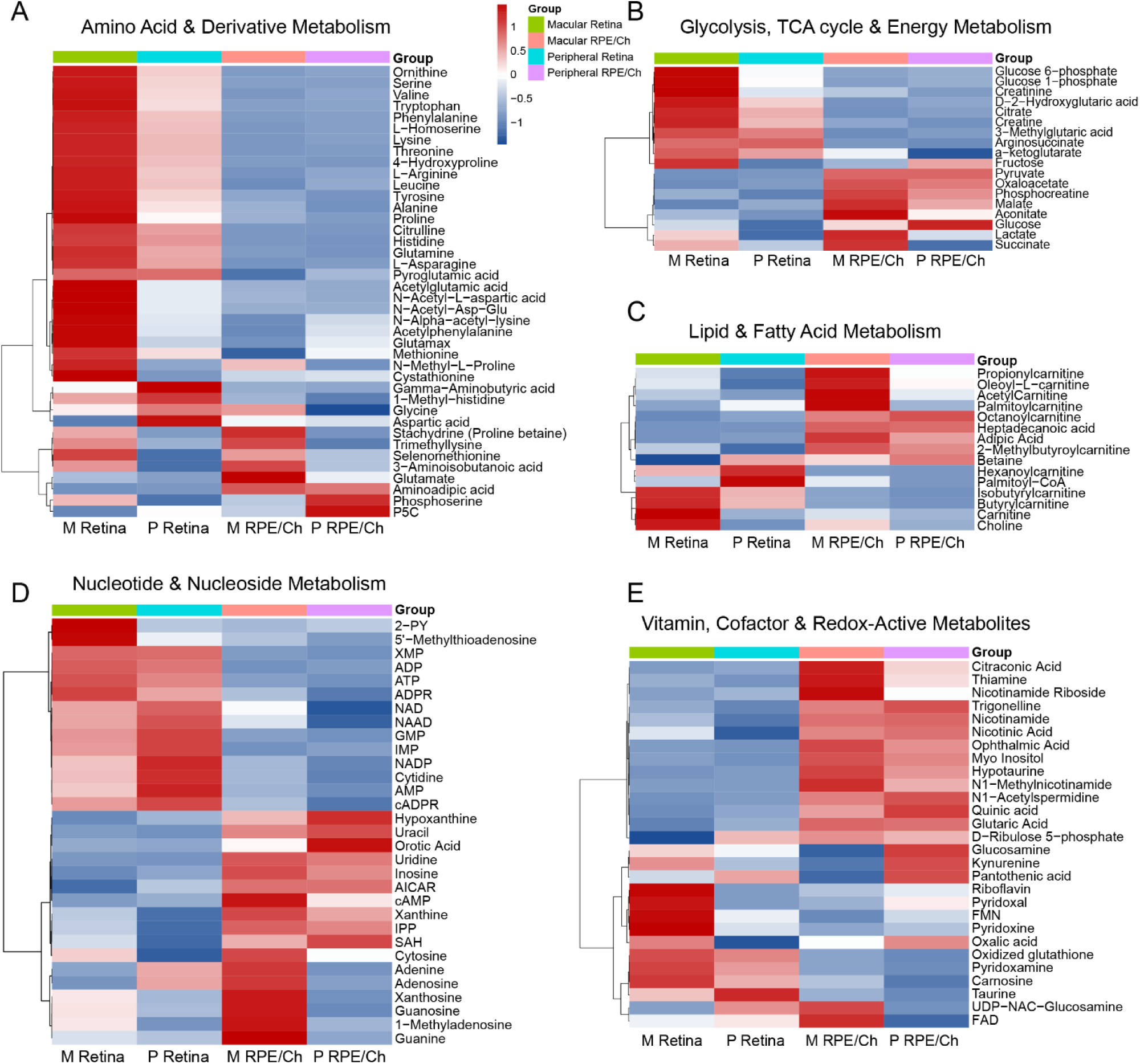
Heatmap showing Z-score normalised metabolite levels in neural retina and RPE/choroid from the macula (M) and periphery (P). Samples include neural macula (M RA), peripheral retina (P RA), macular RPE/choroid (M RPE/Ch) and peripheral RPE/choroid (P RPE/Ch). Each column shows the average for each region and each row represents a metabolite. Red indicates higher levels; blue indicates lower levels. Clustering highlights distinct metabolic patterns between macular and peripheral tissues and between retina and RPE/choroid.

Pyridoxamine, taurine, NADP and oxidised glutathione tended to be higher in the neural retina, whereas nicotinamide riboside, nicotinamide and N1-methylnicotinamide were relatively enriched in the RPE/choroid (**Fig. 5E**).

A schematic summary of these metabolic patterns across the macula and periphery is presented in **Fig. 6**.

**Figure 6.**
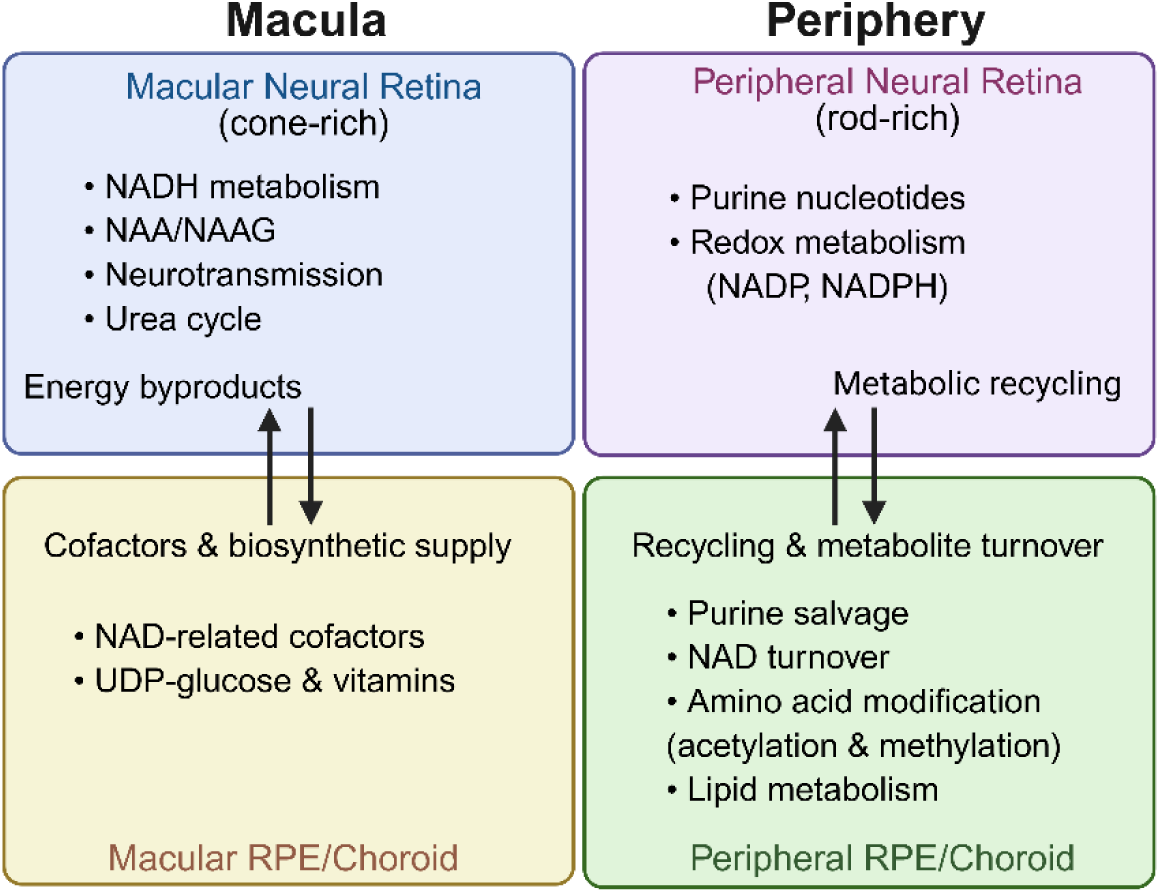
Schematic of complementary metabolic coupling between neural retina and RPE/choroid in macula vs periphery.

## Discussion

This study characterised regional and tissue-specific metabolic differences in the human neural retina and RPE/choroid following short-term *ex vivo* metabolic recovery. Targeted metabolomics revealed that these tissues are not metabolically uniform but show coordinated regional differences. These findings suggest that metabolic interaction between the neural retina and RPE/choroid is adapted to the distinct functional demands of central and peripheral vision.

### Macular region: energy and biosynthetic coordination

Compared with the peripheral neural retina, the neural macula had higher levels of NADH, NAA and NAAG. Cones have more mitochondria than rods, consistent with their greater energetic demands. The increased NADH observed in the neural macula may therefore reflect greater mitochondrial reducing power. The advantage of high reducing power helps preserve glutamate from mitochondrial oxidation (18), which maintains glutamate availability for the synthesis of NAA and NAAG. Excessive reducing power can also promote reductive stress, which is associated with macular degeneration (19). We previously demonstrated that the human neural retina relies heavily on aspartate and glutamate, whereas the neural macula preferentially utilizes pyruvate to support rapid NADH generation (10). Glutamate and aspartate are direct substrates for NAA and NAAG synthesis, while NAAG can be hydrolysed to regenerate these amino acids, forming a reversible metabolic pool. These findings suggest that the neural macula is metabolically adapted to support the exceptionally high energetic and neuronal demands of cone-mediated central vision.

NAAG is one of the most abundant neurotransmitter peptides in the brain, where it regulates glutamate homeostasis through dual mechanisms (11). Acting on astrocytes, NAAG can reduce excessive glutamate release and stimulate the secretion of protective factors (20). Given the functional parallels between astrocytes and Müller glia, similar mechanisms may function in the retina. Elevated NAAG in the neural macula may therefore reflect a dual role: limiting excitotoxic glutamate while also serving as a reservoir to supply glutamate and aspartate in addition to those provided by the RPE (10). Together with the enrichment of NAA, these findings suggest that high neuronal metabolic demand renders the macula more dependent on glio-neuronal metabolic coupling to sustain continuous visual signalling.

We found that the macular RPE/choroid showed higher levels of NAD-related metabolites, including NAD, NADP, NAAD and cADPR, as well as the nucleotide sugar UDP-glucose, compared with the peripheral RPE/choroid. This enrichment highlights enhanced NAD metabolism in the macular RPE/choroid, including NAD biosynthesis, interconversion and turnover. In addition to redox functions, NAD is actively degraded through NAD-consuming enzymes, including poly (ADP-ribose) polymerases (PARPs), sirtuins and CD38 (21). Enrichment of UDP-glucose highlights enhanced nucleotide sugar metabolism in the macular RPE/choroid, where UDP-glucose supports glycogen synthesis and glycosylation reactions essential for membrane integrity and protein modification. The coordinated enrichment of NAD-related metabolites in the macular RPE/choroid together with increased NADH in the neural macula suggests that metabolic coupling between these tissues plays an important role in maintaining redox balance and meeting the high energetic demands of central vision.

### Peripheral region: nucleotide turnover and antioxidant defence

Compared with the neural macula, the rod-rich peripheral retina had higher levels of antioxidant-associated metabolites, including NADPH, hypotaurine and taurine. This suggests that the peripheral retina maintains a strong redox buffering capacity. Compared with the macular RPE/choroid, the peripheral RPE/choroid had higher levels of purine salvage and degradation-related metabolites, including inosine and hypoxanthine, together with downstream NAD-related metabolites such as nicotinamide and N1-methylnicotinamide. This pattern suggests that nucleotide recycling and NAD turnover are more prominent in the peripheral RPE/choroid. Together, these findings indicate complementary metabolic roles in the peripheral region, with the neural retina enriched in antioxidant-related metabolites and the RPE/choroid enriched in recycling and NAD turnover.

Findings in the peripheral RPE/choroid further revealed higher levels of amino acid-related metabolites, including acetylphenylalanine, phosphoserine and arginine, together with the methylation-related metabolite S-adenosylhomocysteine (SAH). These profiles point to active protein modification and amino acid turnover in the peripheral RPE/choroid. This suggests that it is metabolically specialised for sustained substrate recycling to support the long-term activity of photoreceptors in low-light conditions. In addition, the peripheral RPE/choroid had higher levels of N1-methylnicotinamide, a downstream metabolite of NAD metabolism, compared with the neural retina. 1-MNA has been reported to modulate metabolic homeostasis by stabilising SIRT1 protein and thereby influencing hepatic glucose and lipid handling (22). In other contexts, 1-MNA has also been shown to stimulate lipolysis directly (23). These features suggest that the peripheral RPE/choroid contributes to long-term metabolic homeostasis in the rod-rich peripheral retina through amino acid turnover, methylation-related metabolism and NAD turnover.

### Regional metabolic interdependence and disease vulnerability

Our data suggest that metabolic relationships between the neural retina and RPE/choroid differ between the macula and periphery, reflecting the distinct functional demands of central and peripheral vision. In the macula, coordinated glycolytic, TCA cycle and NAD-dependent metabolism support the high energetic requirements of cone-mediated high-acuity vision. This metabolic profile may also increase sensitivity to disturbances in energy production, redox homeostasis or RPE/choroidal support, potentially contributing to the selective vulnerability of the macula in age-related macular degeneration. Understanding these metabolic relationships may provide insights into pathways that maintain retinal homeostasis and guide the development of future metabolic therapies.

Several limitations should be considered when interpreting these findings. The study relies on *ex vivo* human donor tissue, which may be influenced by postmortem delay, donor health status and storage conditions. Targeted metabolomics provides focused coverage of key metabolic pathways but does not capture the full metabolome, particularly lipid classes and signalling molecules that may be relevant to retinal physiology and disease. In addition, metabolite abundances represent steady-state concentrations and cannot distinguish metabolite production from consumption without metabolic flux analysis. Finally, the RPE and choroid were analysed together, limiting interpretation of their individual metabolic contributions.

In conclusion, this study compares metabolic profiles in the human macular and peripheral neural retina and RPE/choroid. Our findings indicate that the neural retina and RPE/choroid have complementary but regionally specialised metabolic roles that support the distinct functional demands of central and peripheral vision. The macula has metabolic features consistent with high energetic demand, providing a potential metabolic basis for its selective vulnerability in macular disease.

## Methods

### Human retinal explant culture

Human donor eyes were used in accordance with approvals granted by the Human Research Ethics Committee of the University of Sydney (Protocol Numbers: 2016/282) and Sydney Local Health District Ethics Review Committee (2020/ETH03339). Retinal dissection followed standard procedures as outlined previously (24). In brief, donor eyes, which had been obtained from the New South Wales Tissue Bank following corneal removal for transplantation, were preserved in CO2-independent medium at 4°C until dissection. The iris, lens and vitreous were removed to expose the posterior eyecup. The neural retina was then carefully detached from the retinal pigment epithelium (RPE) and transferred to a fresh dish. A circular retinal sample, with a diameter of 5 mm, was trephined from either the macula or the mid-peripheral superior retina using a punch biopsy instrument (#BP-50F, Kai Medical). The mid-peripheral region was identified as the midpoint between the fovea and ora serrata. Neural retina and RPE/choroid explants from the macular and peripheral regions were cultured separately in DMEM supplemented with 1× B27, 1% penicillin-streptomycin and 1% FBS. After 4 hours’ incubation period, the neural retina and RPE/choroid tissues were briefly rinsed in ice-cold 0.9% NaCl, snap-frozen in liquid nitrogen and stored at -80°C until metabolite extraction.

### Metabolite extraction and preparation

To characterise regional metabolic differences, we analysed paired macular and peripheral neural retinal and RPE/choroid explants from human donors following 4-hour *ex vivo* metabolic recovery in culture media. Metabolites were extracted using the method previously described (10, 25).

Briefly, retinal or RPE/choroid tissues were homogenised in a cold extraction buffer containing methanol, chloroform and water at a ratio of 800:200:50. The homogenates were incubated on ice with gentle rocking for 20 minutes and then centrifuged at 13,000 × g for 10 minutes at 4°C. The supernatants were collected and freeze-dried for mass spectrometry analysis.

### Targeted Metabolomics

Targeted metabolomics was conducted using liquid chromatography-mass spectrometry (LC-MS) following previously described methods (26, 27). A total of 133 metabolites from major metabolic pathways were quantified (See details in **Supplementary Table 1**). Metabolite analysis was performed using a Shimadzu LC Nexera X2 UHPLC coupled with a QTRAP 5500 LC-MS system (AB Sciex, Framingham, MA). Data analysis was carried out using MultiQuant 3.0.2 (AB Sciex) and Agilent MassHunter Quantitative Analysis Software. Metabolite abundances were normalised to the total protein concentration of protein pellets remaining after metabolite extraction. In this study, metabolites from macular and peripheral retinal and RPE/choroid tissues of eight human donors were analysed.

### PLS-DA, Volcano Plot and Heatmap for Metabolomic Data Analysis

Metabolomic data were analysed using MetaboAnalyst 6.0 (28) (https://www.metaboanalyst.ca). Partial least squares-discriminant analysis (PLS-DA) was performed to visualise metabolic differences between groups, and volcano plots were used to identify differential metabolites.

Metabolites with a fold change ≥ 1.5 and a p-value < 0.05 were considered significantly different. For heatmap analysis, metabolite intensities from neural retina and RPE/choroid from macula and periphery were normalised to total protein, log₂-transformed and Z-score scaled for each metabolite. Heatmaps were created using the online platform bioinformatics.com.cn (29) with hierarchical clustering (Euclidean distance, Ward linkage), where colours indicate relatively low (blue) to high (red) abundance.

### Statistical Analysis

Differential metabolite analysis between macular and peripheral neural retina or RPE/choroid samples was performed using log₂ fold-change calculations. For each metabolite, the mean abundance in the macula was divided by that in the periphery, and the resulting ratio was log₂-transformed. A positive log₂ fold change indicated higher abundance in the macula, whereas a negative value indicated higher abundance in the periphery. Metabolite levels between paired groups were compared using paired t-tests after assessment of data normality. Results are presented as mean ± standard error of the mean (SEM). Statistical significance was defined as *p* < 0.05 for both bar-chart and volcano-plot analyses. All statistical analyses were performed using GraphPad Prism version 11.0.0 (GraphPad Software, San Diego, CA, USA).

## Acknowledgements

This study is supported by the Retina Research Foundation (JD), the Lowy Medical Research Institute (MG, LZ and TZ) and the National Health and Medical Research Council of Australia (Investigator Grant GNT1195021 to MG). We gratefully acknowledge the NSW Eye Bank, the eye donors and their families, whose generous contributions made this study possible.

